# The contribution of Mediterranean connectivity to morphological variability in Iron Age sheep of the Eastern Mediterranean

**DOI:** 10.1101/2022.12.24.521859

**Authors:** Sierra A. Harding, Angelos Hadjikoumis, Shyama Vermeersch, Roee Shafir, Nimrod Marom

## Abstract

The movement of livestock across the Mediterranean is well-documented in the Neolithic era, but its significance during subsequent periods has received less attention. Here we start addressing this lacuna by analyzing astragal bone morphology from four coastal and inland sites in Israel and Cyprus, seeking potential evidence for maritime connections between sheep populations in the Iron Age eastern Mediterranean. Employing an established geometric morphometric protocol, we investigated the hypothesis that intra-site morphological variability is higher in coastal sites, reflecting greater genetic diversity in the livestock populations. While our findings support this hypothesis, the conclusions are constrained by contextual and sample size limitations.

## Introduction

The genetic and phenotypic variability of livestock is a multifaceted phenomenon relating to animal improvement (Aniceti & Albarella, 2022; Davis, 2008; Robin & Clavel, 2018; Valenzuela-Lamas & Albarella, 2017), functional-morphological habitat adaptation (Harbers et al., 2020), inbreeding or divergence between geographically distinct populations (Pöllath et al., 2019; Trentacoste et al., 2018), which is affected in turn by synanthropic mobility (Frantz et al., 2020). Until recently, most studies on livestock mobility have focused on the spread of domesticated animals across space and through time (Daly et al., 2018; Davis & Simões, 2016; Krause-Kyora et al., 2013; Ottoni et al., 2013). Recent research has also focused on later periods, revealing how local livestock improvement, geographical and cultural isolation, human demographic expansion, and long-distance colonization and trade shaped the rich diversity of livestock populations that rapidly disappeared in the modern period (e.g., Fuks & Marom, 2021; Haruda 2017; Meiri et al., 2017; Nieto-Espinet et al., 2020).

By tracing subtle dissimilarities between animals from different times and locales, techniques like ancient DNA and geometric morphometrics allow us to perceive some of the genetic and phenotypic patterns in ancient livestock (Colominas Barberà et al., 2019; Evin et al., 2015; Haruda et al., 2019; Pöllath et al., 2019). In this study, we attempt to isolate and identify the contribution of maritime mobility to livestock biological diversity in Antiquity by investigating the phenotypic variability of domestic sheep from coastal and inland archaeological sites in the eastern Mediterranean (Muñiz et al., 1995; Valenzuela-Lamas et al., 2018).

This study emerges from the current paradigm underpinning the importance of connectivity in Mediterranean history and archaeology. This paradigm was first put forward explicitly by Horden and Purcell (2000), and its application has been extended to the Bronze and Iron Age Iron Age Mediterranean by the magisterial work of Cyprian Broodbank (2013). It states that constant material exchange between communities was a basic tenant of pre-Modern Mediterranean life. This exchange alleviated the uneven distribution of risk and opportunity between different micro-environments in a chronically unpredictable environment. A deeply-ingrained sociocultural adaptation to the *longue durée* features of Mediterranean geography, exchange was often a maritime affair capitalizing on the advantages of the Sea as a medium for rapid transportation.

But were livestock part of this constant maritime exchange in the Bronze and Iron Ages? Mediterranean maritime trade in the Bronze and Iron Ages immediately evokes the riches of the Uluburun shipwreck (Cline & Yasur-Landau, 2007; Pulak, 1998). However, smaller-scale trade is known from the late second millennium BCE onward in the eastern Mediterranean by evidence of the mixed cargoes of amphorae, scrap metal and other mundane items found, for example, in the Cape Gelidonya and Point Iria shipwrecks (Bass, 1967; Bass, 1961; Phelps et al., 1999). It is reasonable to presume that livestock were among the goods transported via *cabotage* or larger-scale maritime trade, especially male breeding stock, although concrete osteoarchaeological evidence from a shipwreck is not known before the end of Classical Antiquity. Complete bones of a single male sheep were recovered from the Ma’agan Mikhael B shipwreck, interpreted to be a merchantman wrecked off the Carmel coast in the middle of the 7^th^ century CE. No butchery or consumption marks were found on the bones, which suggest that they represent the remains of an animal transported alive on board (Harding, 2021).

Earlier evidence for maritime trade in livestock in the east Mediterranean exists. For example, a European haplotype observed in the mitochondrial DNA of south Levantine pigs was traced back to an introduction event at the turn of the second millennium BCE (Meiri et al., 2013). Similarly, patterns of greater body size variability were observed in sheep from Iron Age coastal sites in the Levant in two recent biometric studies (Chahoud et al., 2023; Harding et al., 2023). This enhanced variability could be related to the introduction of non-local breeding stock to coastal settlements via maritime trade networks, as suggested in both studies; however, as their authors add, changes in livestock height and robusticity could also result from changes in nutrition, castration, and other herd maintenance, optimization and management practices.

In this study, we directly address the relationship between morphological variation in livestock and access to maritime trading networks in Antiquity. To do so, we focused on Iron Age sheep remains from two coastal and two inland sites in the southern Levant and Cyprus. This region was a hub of maritime activity during that period, when trans-Mediterranean maritime connections stretched out from the Levantine coast to Gibraltar (Brody, 2002; Broodbank, 2013; Eshel et al., 2018). We expect that this extensive maritime network also includes trade in animals, and therefore these sites provide a suitable setting to explore the link between maritime connectivity and livestock mobility.

We measured sheep morphological variability using a published geometric morphometric protocol that was successfully applied to study morphological variability in the astragali of geographically distinct populations of domestic sheep (or ‘breeds’) (Pöllath et al., 2019). Astragali are considered ideal for the investigation of phenotypic differences between livestock populations (Colominas et al., 2019) because they ossify early (Albarella & Payne, 2005), leaving shorter time for ontogenetic morphological change. In addition, astragal morphology appears to be independent of nutrition and sex (Boessneck 1969; Davis 2000). These characteristics of the astragal, therefore, reduce from the outset the possible effect of sex composition and ontogenetic histories, including nutrition, on its shape. Possible effects of other environmental factors, such as terrain or climate, were controlled by choosing samples obtained from sites in similar topographic and climatic settings. Given this, our (alternate) hypothesis is that the *variability of shape observed in sheep astragali from coastal sites directly exposed to maritime networks will be higher than those from inland sites*.

## Materials and methods

The study comprises of three southern Levant samples from Iron Age 2 Abel Beth Maacah (ABM, N=74), Iron Age 2 Tel Dor (Dor, N=12); pooled Iron Age 2 and Persian Tell Keisan (Keisan, N=14), and a fourth Cypriot sample from Archaic-Classical Lingrin tou Digeni (LTD, N=20). Although every effort was made to achieve a spatiotemporally balanced dataset, the diverse zooarchaeological assemblages included in this study are not homogenous with respect to the sample size from each site and time period. In the case of Dor and Keisan, we analyzed a sample of N<15. Smaller sample sizes have been shown to be less statistically powerful in previous analyses and the results produced here should be interpreted with caution (Cardini et al., 2015; Pöllath et al., 2019). It is also worth noting that the astragali from ABM were recovered from a jar that held more than 400 astragali of diverse artiodactyls (Susnow et al., 2021), while the specimens from Keisan and Dor were derived from refuse layers in a settlement context or, in the case of LTD, from a cultic context. These points will be discussed following the presentation of the results below.

### Study sites

We focus our efforts on the four assemblages chosen because they are in different degrees of remoteness from the sea, and because they have a fair number of astragali from Iron Age contexts, some of which could be clearly identified as sheep. Our study sites are spatiotemporally located in the Iron Age and the Persian period (9th-4th centuries BCE) Eastern Mediterranean (Figure 1):

**Figure 1.**
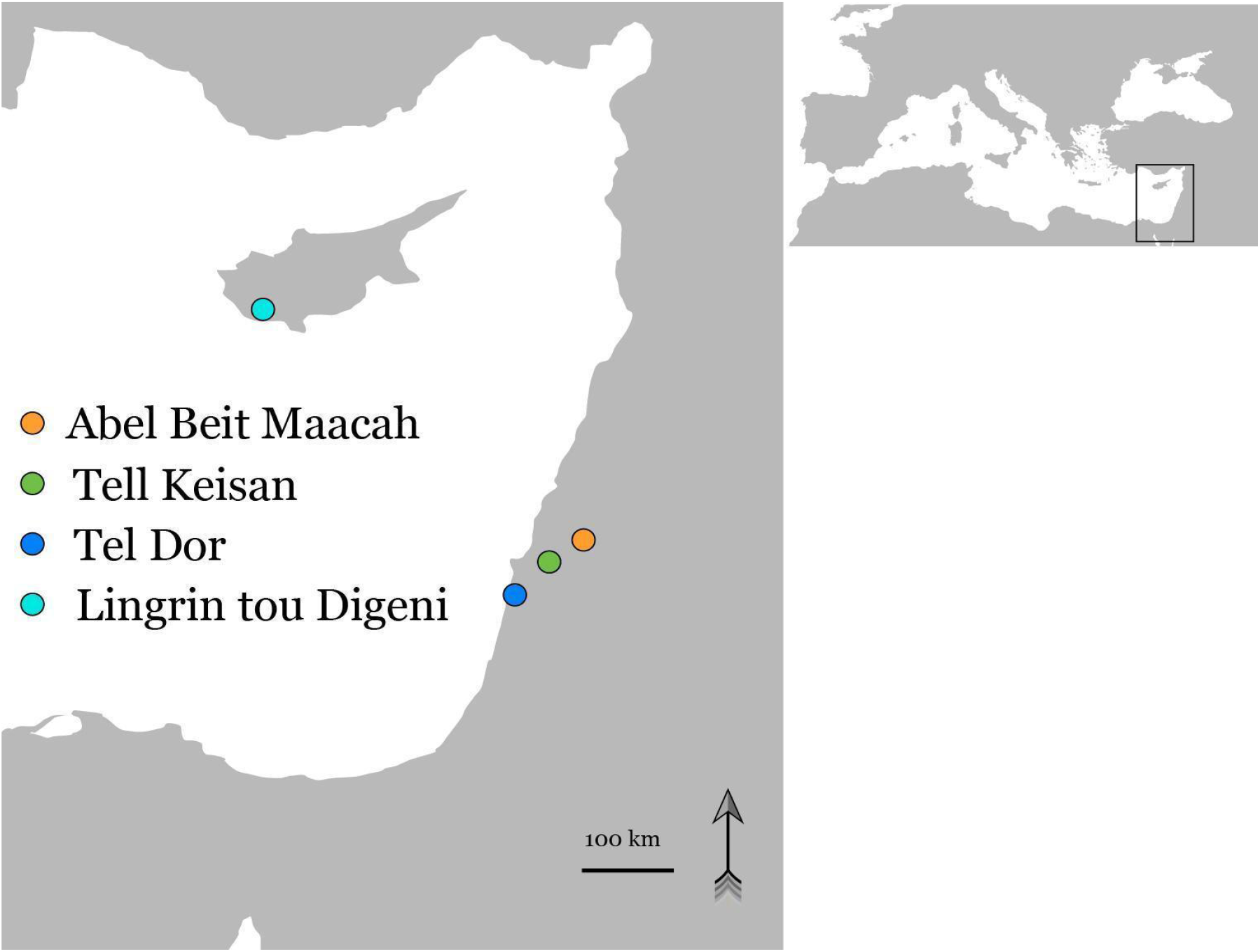
Location map for the sites mentioned in the text, with coastal sites in shades of blue. Base map TheDastanMR, CCo, via Wikimedia Commons, made available under the Creative Commons CCo 1.0 Universal Public Domain Dedication

#### A) Tel Abel Beth Maacah (Inland Settlement)

Tel Abel Beth Maacah (Tel Abil el Qameh), is located at the northern Hula Valley, 37 km east of the coast across the mountainous Galilee and its Mediterranean forest vegetation (Panitz-Cohen and Yahalom-Mack, 2019). Excavations uncovered a sequence from the Bronze Age through the Ottoman period. In the Iron Ages 1 and 2a (IA1–2a, 12^th^–9^th^ centuries BCE), the site was a major urban center (Susnow et al., 2021). The study sample comes from a hoard of astragali found in a jar dating to an IA2a (10^th^–9^th^ centuries BCE) context (Susnow et al., 2021). The settlement’s economy at that period relied mainly on caprines, with a low sheep to goat ratio, with cattle comprising the second most common species (Marom et al., 2020).

#### (B) Tell Keisan (Inland Settlement)

Tell Keisan was settled from the Neolithic to the Hellenistic periods, with peaks in the Middle Bronze Age and the Iron Age (Briend and Humbert, 1980; Stern et al., 1993b). Situated in a valley 8 km east and across the coastal plain from the port city Akko, Keisan served as its agricultural hinterland and a waystation on the road to the hills of the western Galilee (Stern et al., 1993b; Lehmann and Peilstocker, 2012; Aubet, 2014). Tell Keisan had close ties with Tyre throughout the Iron Age 2 and Persian periods, and a strong ‘Phoenician’ influence on its material culture (Lehmann and Peilstocker, 2012). The study sample includes specimens from deposits dating to the Iron Age 2 and the Persian period. The faunal remains are being analyzed by one of us (SV) and represent general discard of animal bone remains comprising mostly caprines.

#### (C) Tel Dor (Coastal Settlement)

Tel Dor (Khirbet el-Burj) is located on the Carmel coast 21 km south of Haifa, in a Mediterranean climate two kilometers from the foot of Mt. Carmel. It was an important seaside settlement from the first half of the second millennium BCE through the first half of the first millennium CE. Dor has been one of the few natural anchorages along the southern Levantine coast, and therefore a major port and maritime activity center. The Iron Age sequence at Dor can be divided into two subsequences: the broadly Phoenician ‘Early Sequence’ (1100–850 BCE), with connections to Egypt, Cyprus, and the Mediterranean; and the ‘Late Sequence’ (850–630 BCE) with an inland-facing Israelite administrative center up until the conquest of the Neo-Assyrian empire in 734/2 BCE (Gilboa et al., 2015; Gilboa and Sharon, 2017). The animal economy at Tel Dor remained remarkably stable through time, with a generalized Mediterranean mixed economy dominated by sheep and goat (Raban-Gerstel et al., 2008; Sapir-Hen et al., 2014; Bartosiewicz and Lisk, 2018). The geometric morphometric sample comes from different Iron Age 2 contexts in Area D5, which represent general discard of animal bone remains.

#### (D) Lingrin tou Digeni (Coastal Settlement)

The site of Kouklia-Lingrin tou Digeni (hereafter ‘Lingrin tou Digeni’) is a sanctuary in Cyprus that dates mainly to the Archaic and Classical periods. It is located near Paphos, less than one kilometer from the coast in a hilly terrain covered by Mediterranean forest. Emergency excavations, which took place after looting was reported in the area, revealed extensive evidence of ritual activity in the form of relevant architectural remains (e.g. courtyard with altar), large numbers of votive human and animal figurines, as well as an associated large faunal assemblage (Raptou 2008; 2010). These ritual deposits yielded the sample used for the geometric morphometric analysis. The study of the fauna is ongoing by one of us (AH). Preliminary observations from approximately 2,000 identified remains suggest that the assemblage is overwhelmingly sheep/goat (∼95%) with cattle comprising the remaining identified animals.

### Specimens

The geometric morphometric analysis (GMM; for code and data files see https://zenodo.org/doi/10.5281/zenodo.10276146) was conducted on samples of sheep astragali that were osteologically mature at time of death, which happens in the first few months of life. We also chose specimens that are free from surface damage, modifications, burning, pathology, or other anomalies that would inhibit comparison using digital landmarks. Skeletal elements were visually identified as *Ovis* sp. using morphological criteria (Boessneck, 1970; Zeder & Lapham, 2010) and the comparative collection at the Laboratory for Archaeozoology at the University of Haifa. The possibility that wild sheep were included in the analysis is very small for the sites in Israel, where no wild sheep existed during the Iron Age. No mouflon horn core, which is the most reliable morphological feature distinguishing domestic sheep from the Cypriot mouflon, has been identified during the ongoing analysis of LTD faunal remains by AH. Moreover, the assemblage consists of only domestic animals, and most were of young age, further indicating a strong preference for domestic rather than feral sheep. While some specimens from LTD could theoretically represent mouflon (feral sheep), given the absence of wild animals at the site and the preference for young age, such a scenario is highly unlikely.

### Digitization

Digital photographs of the anterior surface of each astragalus were taken in ambient lighting with a Nikon D7500 using an AF-S Nikkor 40mm lens by one of us (RS). The camera was stabilized on the photography table using a 90° stable arm. For each photo, the astragalus was placed in a sand-filled box and a bubble level was used to ensure that the surface was horizontal. A 1 cm scale was placed at the level of the photographed surface. The dataset of digital photos was imported into tpsUtil (v.147) (Rohlf, 2015). Digitization of landmarks and semi-sliding landmarks (collectively ‘landmarks’ below, for brevity) was conducted by laboratory assistant Daria Lokshin Gorzilov, under the direct supervision of SH, using tpsDig232 (v.231) (Rohlf, 2017). We preferred left-side elements; images of right-side elements were flipped in tpsDig 232. The landmark configuration followed the protocol outlined in Pöllath et al. (2019): 11 fixed landmarks (LM) and 14 semi-sliding landmarks (SSLM) were placed around the outline of the dorsal view (Figure 2). A scale was set for each photo within tpsDig.

**Figure 2.**
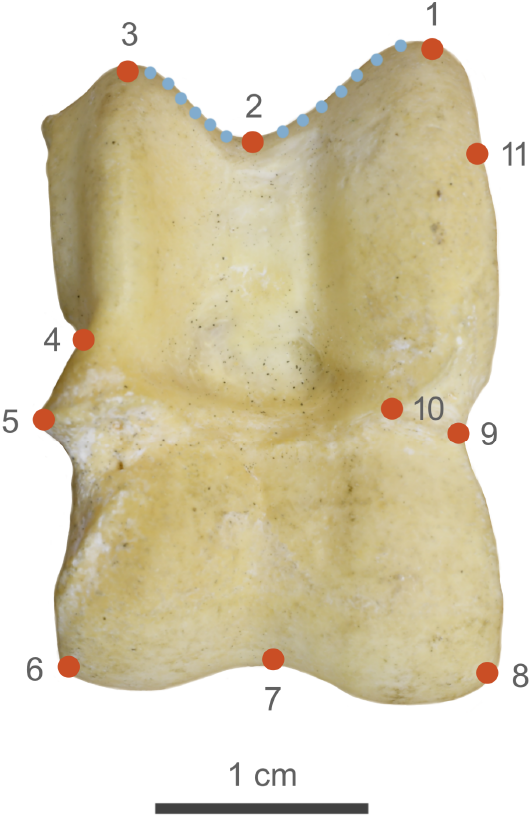
Landmark (red) and sliding semi-landmark (blue) placement, after Pöllath et al. (2019).

### Generalized Procrustes analysis

A Generalized Procrustes Analysis (GPA) was performed using the geomorph::gpagen function (Adams et al., 2022; Adams & Otárola-Castillo, 2013) on the landmark coordinates obtained in tpsDig which produced Procrustes shape variables (Bookstein, 1991). Semi-sliding landmarks were allowed to slide relative to each other during the GPA to minimize the sum Procrustes distances between each specimen and the mean shape (Bookstein, 1996). Centroid sizes, which are calculated as the square root of the sum of the squared distances of the landmarks from the centroid of the specimens, were also produced by the same function. The mean Procrustes shape for each site was obtained from the geomorph::gpagen function for visualization.

### Digitization error

Digitization percentage errors were checked before statistical analysis by using tpsSmall64 (v.1.0) and tpsrelw32 (v.1.53) (Rohlf, 2015) according to the protocol described by Hulme-Beaman (2014: 153). A subset (N=28) of specimens from the dataset were digitized three unique times to test for digitization error; these were randomly selected but as evenly distributed between the study sites as possible. This analysis employs Procrustes distances to quantify both biological variation and digitization error. Firstly, it calculates the Procrustes distance between all specimens to assess the extent of inherent biological variability. Secondly, it computes the Procrustes distance between replicate measurements of each individual, isolating the variability introduced by the digitization process. Finally, by expressing the digitization error as a percentage of the total observed variability, the impact of measurement imprecision on the overall results is evaluated.

### Procrustes analysis of variance (ANOVA)

The possible effect of group structure on the shape coordinates is key to our research hypothesis that morphology varies significantly between sites. Also, we needed to test the allometric effect of specimen size on shape. The effect of both these explanatory variables and the interaction between them on shape was estimated using the residual randomization permutation procedure ANOVA method using the geomorph::procDlm function. The Procrustes ANOVA is used to analyze the effect of different explanatory variables on the Procrustes distances between specimens. The procedure uses a randomized residual permutation procedure to overcome potential breakdown of assumptions related to the normal distribution of residuals made in linear models (Collyer & Adams, 2018).

### Ordination and disparity analyses

We used principal component analysis (PCA, implemented using ‘geomorph::gm.prcmp’) to reduce the dimensionality of the Procrustes transformed landmarks, while retaining most of the variability present among the Procrustes shapes. The first ten principle components were used in a canonical variates analysis (‘Morpho::CVA’) – a standard technique that maximizes the distance between groups – to visualize group structure and measure inter-group Mahalanobis distances in the resulting ordination space. We also conducted a Procrustes disparity analysis on the Procrustes transformed coordinates using geomorph::morphol.disparity. Morphological disparity is estimated as the Procrustes variance for groups, using residuals of a linear model fit. P-values can be obtained using a random residual permutation procedure. Procrustes variance is the trace of the group covariance matrix divided by the number of observations in the group (Zelditch et al., 2014: p. 487). Between-group Procrustes distances were also visualized using neighbor-joining trees (using ape::nj).

### R libraries

GMM analysis and visualization were conducted in R (v.4.1.0) (R Core Team, 2022) using the following packages: ‘geomorph’ (Adams et al., 2022; Adams & Otárola-Castillo, 2013), ‘Morpho’ (Schlager, 2017), ‘tidyverse’ (Wickham et al., 2019), and ‘ggsci’ (Xiao, 2018).

## Results

### Digitization error

The mean Procrustes distance between re-digitized specimens was 0.034, and among the full dataset, 0.118; the percent of digitization error is therefore estimated at 29.37%. This percent of digitization error is within the range observed in similar intraspecies GMM studies (Harding, 2021, p. 59; Hulme-Beaman, 2014, pp. 164–167 and table 6.2).

### Centroid size and allometric pattern

Median centroid sizes are plotted in Figure 3A and are significantly different (P<0.001; see ANOVA table in Supplement 1a) between sites. The allometric effect of size on shape data is statistically significant (P = 0.009) but has a negligible effect (R^2^ = 0.01) in comparison to that of group (P = 0.001, R^2^ = 0.11). The effect of the interaction between group and size on shape is not significant (P = 0.068). The full model output appears in Supplement 1 (10.5281/zenodo.10276840).

**Figure 3.**
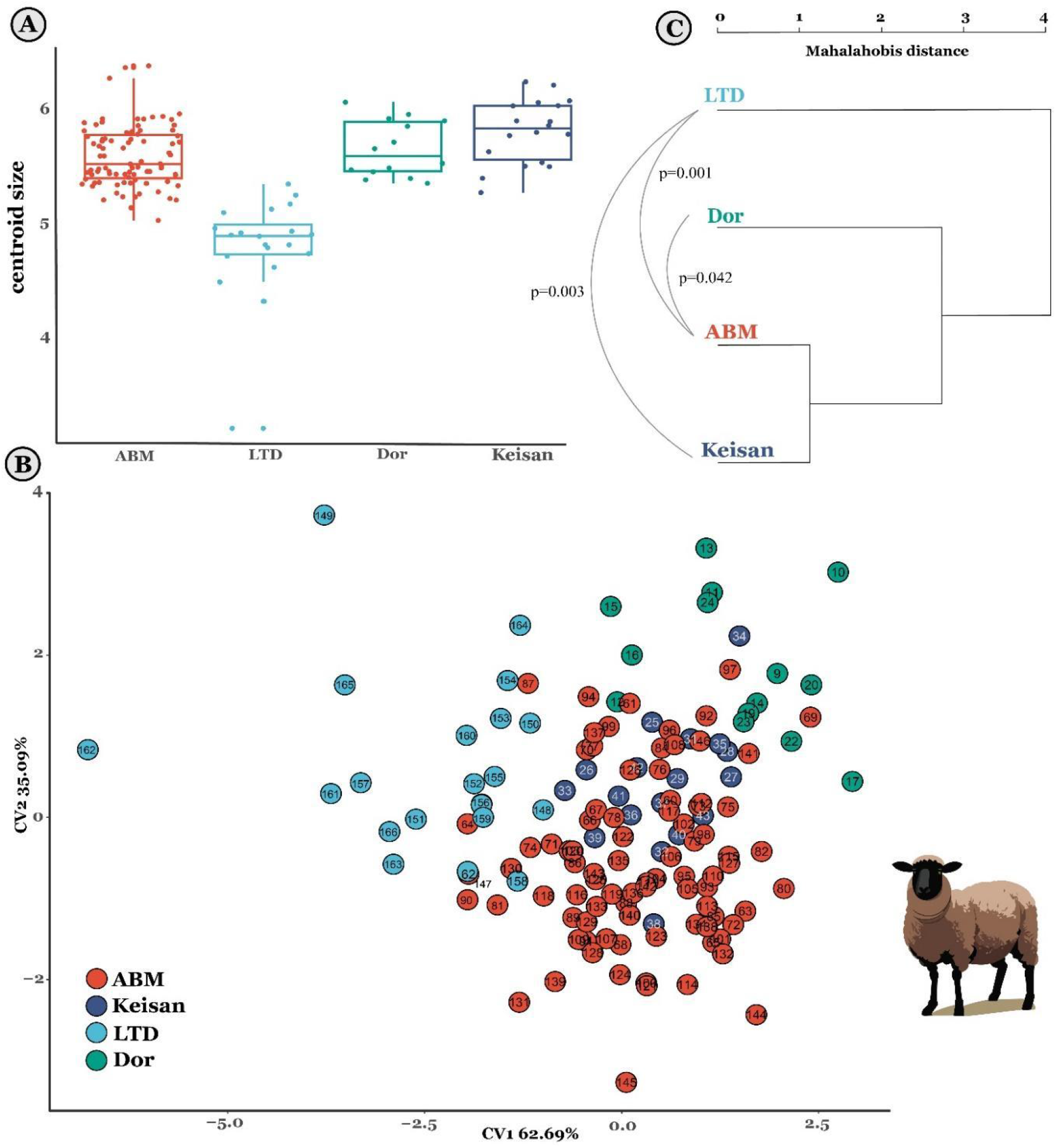
Boxplots present specimen centroid sizes (A) of astragali from Iron Age 2 Tel Abel Beth Maacha (ABM; n=87), Iron Age 2 Tel Dor (n=14), and the combined Iron Age 2 and Persian period samples from Tel Keisan (n=19). (B) Canonical variate ordination of the first ten principal components obtained from a shape PCA of the Procrustes transformed landmark data. (C) Dendrogram of the Mahalanobis distance between groups, calculated from the CVA ordination data. P-values represent significant inter-group pairwise differences in the Procrustes variances. Specimen numbers refer to the specimen info file at https://zenodo.org/doi/10.5281/zenodo.10276146

### Ordination

The first two canonical variates capture most of the variability in the dimensionality-reduced dataset of the first 10 PCs (∼98%)(Figure 3B). The ABM and Keisan specimens cluster together near the origin, with the Dor and LTD data occupying higher and lower regions along CV1, respectively, and a higher position on CV2. The Mahalanobis distance between the groups, when presented in a dendrogram (Figure 3C), reveals a geographic structure similar to the astragal shape data. The classification output of the CVA analysis is in Supplement 1 (10.5281/zenodo.10276840).

### Disparity

The Procrustes distances between groups, calculated from the landmark configurations, are presented in Table 2, and as an unrooted neighbor-joining tree in Figure 4A. The tree suggests a similar geographic structure to that obtained by the ordinated CVA results. The similarity between the results of two different analyses—direct measurements of disparity on landmark data (Table 2) and on eigen-ordinated coordinates (Figure 3) — support the statistical results. A random permutation procedure produced statistically significant differences among the Procrustes distances calculated between the easternmost inland site, ABM, and the coastal samples. Finally, Figure 4C presents the intra-group variance in the Procrustes transformed landmark data in each group in relation to its sample size. Notably, the coastal assemblages have a much higher intra-sample variance than inland ABM; after normalization, the amount of variance per site can be observed to be independent of sample size.

**Table 1.** Site names, codes, dates, and sample sizes.

| <u>Site Name</u> | <u>Site Code</u> | <u>Zone</u> | <u>Dating</u> | <u>Sample N=</u> |
| --- | --- | --- | --- | --- |
| Abel Beth Maacah | ABM | Inland | Iron Age II<br>(10<sup>th</sup>–9<sup>th</sup> c. BCE) | 74 |
| Tell Keisan | Keisan | Inland | Iron Age II–Persian Period<br>(950–332 BCE) | 14 |
| Tel Dor | Dor | Coastal | Iron Age II<br>(950–539 BCE) | 12 |
| Lingrin tou Digeni | LTD | Coastal | Archaic–Classical Period<br>(700–332 BCE) | 20 |

**Table 2.** Pairwise absolute differences between sample variances (below diagonal) and p-values (above diagonal) obtained from a randomized residual permutation procedure (N_permutations_ = 1000).

|  | ABM | Keisan | Dor | LTD |
| --- | --- | --- | --- | --- |
| ABM |  | 0.96 | 0.04 | 0.001 |
| Keisan | 0.0000507 |  | 0.08 | 0.003 |
| Dor | 0.0020000 | 0.0020517 |  | 0.148 |
| LTD | 0.0037418 | 0.0037926 | 0.0017409 |  |

**Figure 4:**
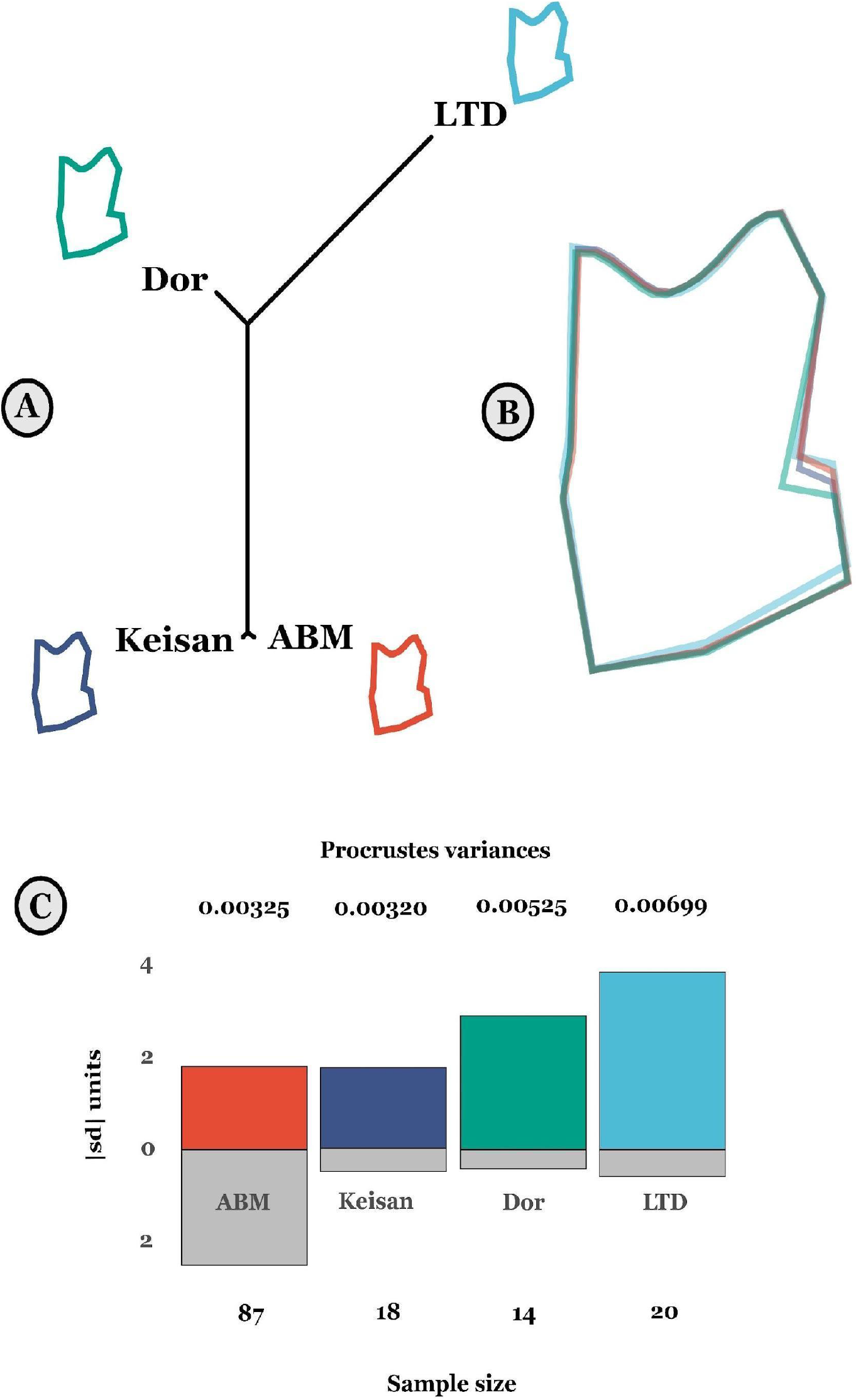
(A) Procrustes distances between groups presented as a neighbor-joining tree; mean shape for each group plotted separately at each tip. (B) Superimposition of the mean shapes for each group. (C) Standardized absolute Procrustes variances within each site in relation to sample size. Absolute values of the standardized columns appear above (distances) and below (sample size) their respective columns.

## Discussion

Our analysis returned results related to size and shape variability within and between sites. First, with respect to size, we observe a significant difference between sites, with the Cypriot LTD showing notably smaller size than the other, southern Levantine sites. The astragalus geometric morphometric results suggest that slightly larger sheep were present in Tell Keisan, which is like the result obtained by a biometric study of the sheep remains from the site (Harding et al., 2023). The smaller size of the Dor specimens also resonates well with the results from other Levantine coastal sites in the Iron Age (Chahoud et al., 2023). The specimens from Cyprus are especially small. We do not know how universal this pattern may be on the island during the Iron Age, and this observation requires further investigation.

Second, morphological variability appears to be greater in coastal sites compared to inland sites, suggesting a morphological cline that to the best of our knowledge has not been noted before. This observation does not appear to be related to sample size, as the site with the largest sample size, ABM (N=87), has approximately half the morphological variance of LTD (N=20). Third, a clear east-west morphocline emerges: morphological similarity between sites maintains the rank order of the geographical distance between them. Significance values mark a clear similarity between Dor/LTD, the coastal sites, and ABM, the easternmost site. Such morphological clines were noted among various mammalian taxa in the past (e.g., Evin et al. 2013; Ottoni et al., 2013; Price et al., 2023; Slim & Çakırlar, 2023).

The interpretation of the data could be straightforward: Samples from coastal sites reflect a morphologically variable population of sheep in the Iron Age. The importance of the sea as a connective medium is apparent in that the sheep from Dor are morphologically closer to the sample from Cyprus than to inland sheep from the nearby Galilee at Tell Keisan. Higher shape diversity observed in the Mediterranean coastal sites strongly suggests our samples from these sheep populations had more exposure to non-local animals, which would support our hypothesis. This general phenomenon of anthropogenic increase in alpha diversity from maritime transportation of animals is a well-known from human colonization of islands (e.g., Cucchi & Vigne, 2006; Hulme-Beaman et al., 2018; Vigne et al., 2014), and from the spread of phenotypes of already domesticated animals, as with the non-agouti cats spread along sea lanes in the historical past (Todd, 1977).

This interpretation, however, should be discussed in the context of the spatiotemporal resolution of our study. Specialists in the archaeology of the Iron Age Levant are used to fine contextual resolutions and a high, almost decadal-scale temporal record obtained from radiocarbon dating and very refined ceramic chronologies (Finkelstein & Piasetzky, 2011; Gilboa & Sharon, 2003; Mazar, 2011). To them, the approximately five centuries spanned by our sample and the site level contextual resolution may seem too coarse to be meaningful. We can answer such concerns in two ways. The first is by noting the trivial fact that we must pool chronostratigraphic phases to even remotely approach a suitable sample size of complete sheep astragali from each site. The second, which is less trivial, is that we should not assume that the pace of spread of livestock morphotypes is related to the pace of change in ceramic assemblages on which archaeological chronostratigraphy is ultimately based. Cultural evolution is rapid in comparison to biological evolution. We therefore think that the scale of time represented by our samples, which spans a few centuries after the Iron Age explosion of trans-Mediterranean trade, is essentially right for observing the more ‘viscous’ spread of sheep morphotypes across large regions.

Confidence in our conclusions is also compromised by issues of zooarchaeological context: we are not sure that the different samples we obtained represent the variability in the geographical region from which they derive in the same way. The large sample from ABM was obtained from a single jar, used as a ‘favissa’ to astragali of different taxa. Even if we accept that the astragali represent random mortality from herds living in the agricultural hinterland of the site, the sample is extremely confined in the temporal sense, because it represents the total collection time of a complete jar, which we estimate in years. This stands in contrast with the longer, decadal or centennial scale accumulations of refuse buried in Keisan and Dor which sample a larger span of time. Regarding spatial variability, in LTD, the high intra-assemblage variation may be due to ritual activities. These work as a sink for livestock variability, as people came from variable distances to make offerings.

An additional factor that may affect ovine astragalus morphotypical diversity is the landscape to which domestic sheep may be adapted to over generations, *i*.*e*., functional mobility adapted to a rough, hilly terrain (closed environment) versus flat plains (open environment) may be reflected in astragalus morphology (Haruda et al., 2019). In the present study, ABM would be considered the most topographically complex, being a high-elevation settlement in the Galilee mountain range (414 m above mean sea level [AMSL]). Dor is located on a coastal promontory (3 m AMSL), and Keisan is situated in a valley near the Galilee foothills (31 m AMSL). Although coastal in terms of linear distance, LTD sits 85 m AMSL in a hilly area of the coastal plain. If functional mobility was a strong factor affecting astragalus morphology, we would expect to see that signal clearly reflected in the data, *e*.*g*., no significant difference between ABM and LTD or between Dor and Keisan. Our results show a significant pairwise difference between ABM and LTD (p=0.001), but not between Dor and Keisan. A previous study echoes this finding, which analyzed astragalus proportions of inland/rough terrain and coastal/flat terrain domestic sheep in the southern Levant using an astragalus dimension index (lateral depth/greatest lateral length; distal breadth/lateral depth) (Harding et al., 2023). The proportions of astragali tended to reflect the environment from which the samples came, yet there was significant overlap in the center of the distribution which muddied any stark distinction between open and closed landscapes (Harding et al., 2023: 8, Fig. 5). Based on the current findings and this previous study, we do not find sufficient evidence to support the hypothesis that terrain-adapted functional morphology overwhelmingly accounts for the variability that we observe in the present study.

**Figure 5.**
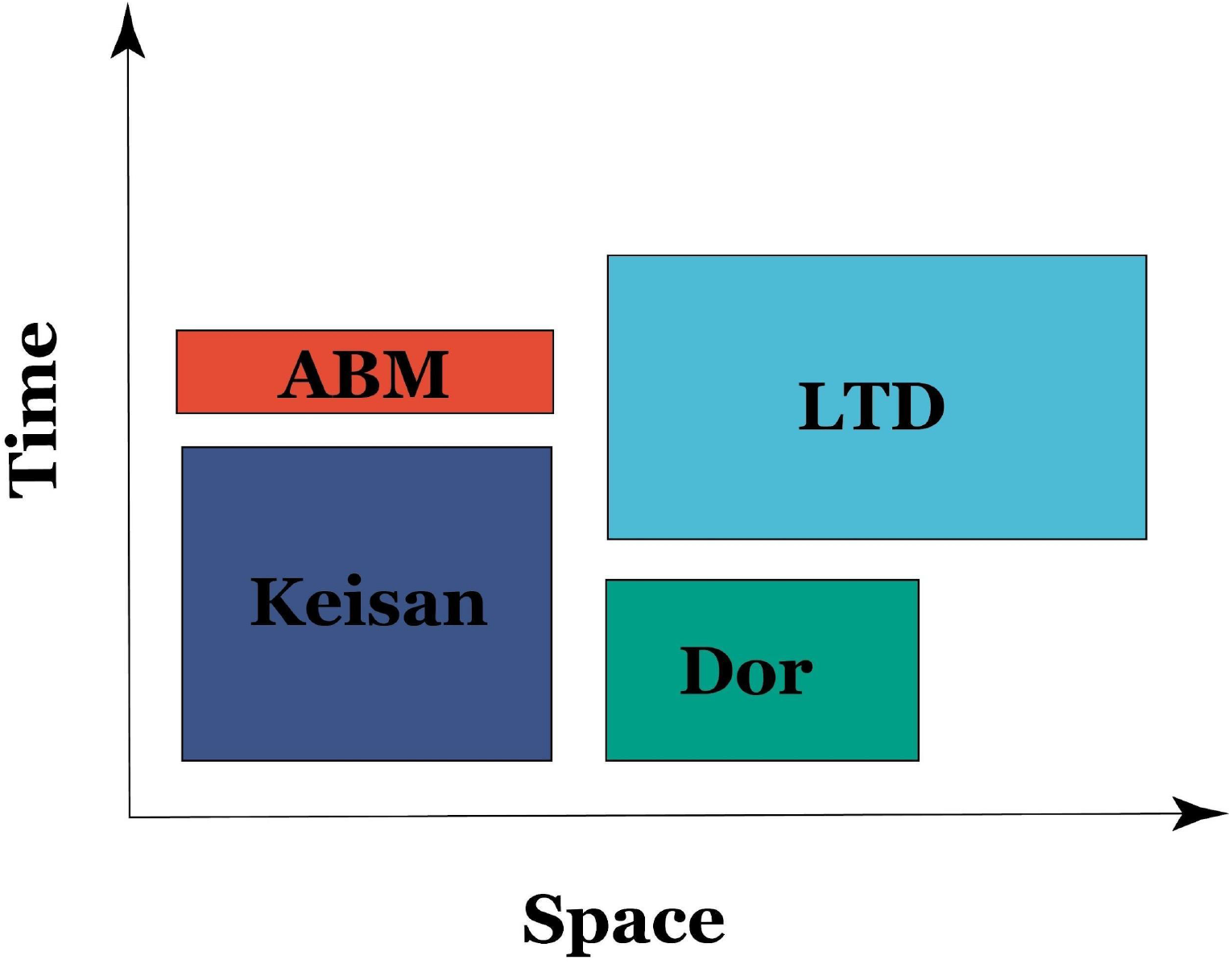
The effect of zooarchaeological context on intra-site shape variability when sample sizes are similar. Rectangle areas, representing intra-site morphological variability, are a function of the spatial variability (x-axis) of the region from which the animals were obtained (large for the ritual site of LTD) and the time period (y-axis) over which they accumulated (very short in ABM).

These effects of zooarchaeological context on intra-site shape variability when sample sizes are similar are illustrated in Figure 5, where rectangle areas, representing intra-site morphological variability, change in proportion to the dimensions of the region from which the animals were obtained and the time period over which they accumulated. Such considerations complicate interpretation, especially on the intra-site variance comparisons. We tend to think that inter-site patterns of variation, especially the inland versus coastal morphotypes, should be more robust.

At this point, our results support the idea that maritime connectivity is related to morphological variability in Iron Age domestic sheep, but with the important caveats mentioned above. The way forward would be to enlarge the number of sampled sites focusing on material recovered from accumulated deposits, thereby eliminating, or at least significantly reducing, idiosyncrasies related to zooarchaeological context. Ancient DNA and stable isotope analyses could help clarify these issues.

## Acknowledgements

We wish to thank Gunnar Lehmann, Ayelet Gilboa, the late Ilan Sharon and Naama Yahalom Mack for allowing us to work on the materials from Tel Keisan, Tel Dor, and Abel Beth Maacah. We also thank Daria Lokshin Gorzilov for her technical assistance with specimen digitization. We would like to thank the Department of Antiquities of the Republic of Cyprus and the excavator Dr. Efstathios Raptou for granting permission to study a sample of astragali from Lingrin tou Digheni. We also wish to thank Louise Le Meillour and two anonymous reviewers for their great help in improving this manuscript.

## Data availability

R codes, TPS files and a specimen catalogue are available at https://zenodo.org/doi/10.5281/zenodo.10276146

## References

Adams, D. C., Collyer, M. L., Kaliontzopoulou, A., & Baken, E. K. (2022). Geomorph: Software for geometric morphometric analyses (version 4.0.4.) [Linux]. https://cran.r-project.org/package=geomorph.

Adams, D. C., & Otárola-Castillo, E. (2013). Geomorph: An R package for the collection and analysis of geometric morphometric shape data. Methods in Ecology and Evolution / British Ecological Society, 4, 393–399.

Aniceti, V., & Albarella, U. (2022). The role of sheep husbandry during the Arab agricultural revolution in medieval Sicily (7th-14th c. AD). Journal of Archaeological Science: Reports, 44, 103529.

Aubet, M. E. (2014). Phoenicia During the Iron Age II Period. In A. E. Killebrew & M. Steiner (Eds.), The Oxford Handbook of the Archaeology of the Levant: c. 8000–332 BCE (pp. 1–13). Oxford: Oxford University Press. 10.1093/oxfordhb/9780199212972.013.046

Bartosiewicz, L., & Lisk, E. (2018). Mammalian Remains. In A. Gilboa, I. Sharon, J. R. Zorn, & S. Matskevich (Eds.), Excavations at Dor, Final Report, Volume IIB—Area G, the Late Bronze and Iron Ages: Pottery, Artifacts, Ecofacts, and Other Studies. Qedem Reports.

Bass, G. F. (Ed.). (1967). Cape Gelidonya: A Bronze Age Shipwreck. American Philopsophical Society.

Bass, G. F. (1961). The Cape Gelidonya Wreck: Preliminary Report. American Journal of Archaeology, 65(3), 267–276.

Boessneck, J. (1970). Osteological differences between sheep (Ovis ariea Linné) and goats (Capra hircus Linné). In D. Brothwell & E. Higgs (Eds.), Science in Archaeology (pp. 331–358). Praeger.

Bookstein, F. L. (1991). Morphometric Tools for Landmark Data. Cambridge University Press.

Bookstein, Fred L. (1996). Landmark methods for forms without landmarks: localizing group differences in outline shape. Proceedings of the Workshop on Mathematical Methods in Biomedical Image Analysis, 279–289.

Briend, J., & Humbert, J.-B. (Eds.). (1980). Tell Keisan (1971–1976): une cité phénicienne en Galilée. Fribourg, Switzerland/Göttingen, Germany: Éditions Universitaires/Vandenhoeck Ruprecht.

Brody, A. (2002). From the Hills of Adonis through the Pillars of Hercules: Recent Advances in the Archaeology of Canaan and Phoenicia. Near Eastern Archaeology, 65(1), 69–80.

Broodbank, C. (2013). The Making of the Middle Sea: A History of the Mediterranean from the Beginning to the Emergence of the Classical World. Thames & Hudson.

Cardini, A., Seetah, K., & Barker, G. (2015). How many specimens do I need? Sampling error in geometric morphometrics: testing the sensitivity of means and variances in simple randomized selection experiments. Zoomorphology, 134(2), 149–163.

Chahoud, J., Vila, E., & Albesso, M. (2023). Livestock management in the Northern Levant during the first millennium BCE. Quaternary International, 662, 63–81.

Cline, E. H., & Yasur-Landau, A. (2007). Musings from a Distant Shore: The Nature and Destination of the Uluburun Ship and its Cargo. Tel Aviver Jahrbuch Fur Deutsche Geschichte / Herausgegeben Vom Institut Fur Deutsche Geschichte, 34(2), 125–141.

Collyer, M. L., & Adams, D. C. (2018). RRPP: An r package for fitting linear models to high-dimensional data using residual randomization. Methods in Ecology and Evolution / British Ecological Society, 9(7), 1772–1779.

Colominas Barberà, L., Evin, A., Campmajó, P., Casas, J., Castanyer i Masoliver, P., Carreras Monfort, C., Guardia, J., Olesti i Vila, O., Pons i Brun, E., Tremoleda i Trilla, J., & Others. (2019). Behind the steps of ancient sheep mobility in Iberia: new insights from a geometric morphometric approach. Archaeological and Anthropological Sciences, 11(9), 4971–4982.

Cucchi, T., & Vigne, J.-D. (2006). Origin and Diffusion of the House Mouse in the Mediterranean. Human Evolution, 21(2), 95.

Daly, K. G., Maisano Delser, P., Mullin, V. E., Scheu, A., Mattiangeli, V., Teasdale, M. D., Hare, A. J., Burger, J., Verdugo, M. P., Collins, M. J., Kehati, R., Erek, C. M., Bar-Oz, G., Pompanon, F., Cumer, T., Çakirlar, C., Mohaseb, A. F., Decruyenaere, D., Davoudi, H., Bradley, D. G. (2018). Ancient goat genomes reveal mosaic domestication in the Fertile Crescent. Science, 361(6397), 85–88.

Davis, S. J. M. (2000). The Effect of Castration and Age on the Development of the Shetland Sheep Skeleton and a Metric Comparison Between Bones of Males, Females and Castrates. Journal of Archaeological Science, 27(5), 373–390.

Davis, S. J. M. (2008). Zooarchaeological evidence for Moslem and Christian improvements of sheep and cattle in Portugal. Journal of Archaeological Science, 35(4), 991–1010.

Davis, S., & Simões, T. (2016). The velocity of Ovis in prehistoric times: the sheep bones from Early Neolithic Lameiras, Sintra, Portugal. O Neolítico Em Portugal Antes Do Horizonte 2020: Perspectivas Em Debate, 2, 51–66.

Eshel, T., Yahalom-Mack, N., Shalev, S., Tirosh, O., Erel, Y., & Gilboa, A. (2018). Four Iron Age Silver Hoards from Southern Phoenicia: From Bundles to Hacksilber. Bulletin of the American Schools of Oriental Research. American Schools of Oriental Research, 379, 197–228.

Evin, A., Cucchi, T., Cardini, A., Strand Vidarsdottir, U., Larson, G., & Dobney, K. (2013). The long and winding road: identifying pig domestication through molar size and shape. Journal of Archaeological Science, 40(1), 735–743.

Evin, A., Flink, L. G., Bălăsescu, A., Popovici, D., Andreescu, R., Bailey, D., Mirea, P., Lazăr, C., Boroneant, A., Bonsall, C., Vidarsdottir, U. S., Brehard, S., Tresset, A., Cucchi, T., Larson, G., & Dobney, K. (2015). Unravelling the complexity of domestication: a case study using morphometrics and ancient DNA analyses of archaeological pigs from Romania. Philosophical Transactions of the Royal Society of London. Series B, Biological Sciences, 370(1660), 20130616.

Frantz, L. A. F., Bradley, D. G., Larson, G., & Orlando, L. (2020). Animal domestication in the era of ancient genomics. Nature Review of Genetics, 21(8), 449–460.

Finkelstein, I., & Piasetzky, E. (2011). The Iron Age Chronology Debate: Is the Gap Narrowing? Near Eastern Archaeology, 74(1), 50–54.

Fuks, D., & Marom, N. (2021). Sheep and wheat domestication in southwest Asia: a meta-trajectory of intensification and loss. Animal Frontiers, 11(3), 20–29.

Gambash, G. (2015). Maritime Activity in the Ancient Southern Levant the Case of Late Antique Dor. ARAM Society for Syro-Mesopotamian Studies, 27(1&2), 61–74.

Gilboa, A., & Sharon, I. (2003). An Archaeological Contribution to the Early Iron Age Chronological Debate: Alternative Chronologies for Phoenicia and Their Effects on the Levant, Cyprus, and Greece. Bulletin of the American Schools of Oriental Research, 332(1), 7–80.

Harbers, H., Neaux, D., Ortiz, K., Blanc, B., Laurens, F., Baly, I., Callou, C., Schafberg, R., Haruda, A., Lecompte, F., Casabianca, F., Studer, J., Renaud, S., Cornette, R., Locatelli, Y., Vigne, J.-D., Herrel, A., & Cucchi, T. (2020). The mark of captivity: plastic responses in the ankle bone of a wild ungulate (Sus scrofa). R. Soc. Open Sci., 7(3), 192039.

Harding, S. A. (2021). Analysis of the Faunal Remains from the Ma‘agan Mikhael B Shipwreck (N. Marom, Ed.) [MA]. University of Haifa.

Harding, S., Vermeersch, S., Ujma, C., Deonarain, G., Susnow, M., Shafir, R., Gilboa, A., Lehmann, G., & Marom, N. (2023). Hoofprints in the sand: A study on domestic sheep (Ovis aries) from Iron Age southern Phoenicia using traditional biometric methods. Quaternary International, 662, 82–93. 10.1016/j.quaint.2023.02.014

Haruda, A. F. (2017). Separating Sheep (Ovis aries L.) and Goats (Capra hircus L.) Using Geometric Morphometric Methods: An Investigation of Astragalus Morphology from Late and Final Bronze Age Central Asian Contexts. International Journal of Osteoarchaeology, 27(4), 551–562. 10.1002/oa.2576

Haruda, A. F., Varfolomeev, V., Goriachev, A., Yermolayeva, A., & Outram, A. K. (2019). A new zooarchaeological application for geometric morphometric methods: Distinguishing Ovis aries morphotypes to address connectivity and mobility of prehistoric Central Asian pastoralists. Journal of Archaeological Science, 107, 50–57.

Horden, P., & Purcell, N. (2000). The Corrupting Sea: A Study of Mediterranean History. Blackwell.

Hulme-Beaman, A. (2014). Exploring the human-mediated dispersal of commensal small mammals using dental morphology: rattus exulans and rattus rattus [Paris, Muséum national d’histoire naturelle]. http://www.theses.fr/2014MNHN0031

Hulme-Beaman, A., Cucchi, T., Evin, A., Searle, J. B., & Dobney, K. (2018). Exploring Rattus praetor (Rodentia, Muridae) as a possible species complex using geometric morphometrics on dental morphology. Mammalian Biology (formerly Zeitschrift Fur Saugetierkunde), 92, 62–67.

Krause-Kyora, B., Makarewicz, C., Evin, A., Flink, L. G., Dobney, K., Larson, G., Hartz, S., Schreiber, S., von Carnap-Bornheim, C., von Wurmb-Schwark, N., & Nebel, A. (2013). Use of domesticated pigs by Mesolithic hunter-gatherers in northwestern Europe. Nature Communications, 4, 2348.

Lehmann, G. (2001). Phoenicians in the Western Galilee: First Results of an Archaeological Survey in the Hinterland of Akko. In A. Mazar (Ed.), Studies in the Archaeology of the Iron Age in Israel and Jordan (pp. 65–112). Sheffield: Sheffield Academic Press.

Lehmann, G. (2021). The Emergence of Early Phoenicia. Jerusalem Journal of Archaeology, 1, 272–324. 10.52486/01.00001.11

Lehmann, G., & Peilstocker, M. (2012). Map of Ahihud (20).

Mazar, A. (2011). Iron Age Chronology Debate: Is the Gap Narrowing? Near Eastern Archaeology, 74(2), 105–111.

Meiri, M., Stockhammer, P. W., Marom, N., Bar-Oz, G., Sapir-Hen, L., Morgenstern, P., MacHeridis, S., Rosen, B., Huchon, D., Maran, J., & Finkelstein, I. (2017). Eastern Mediterranean Mobility in the Bronze and Early Iron Ages: Inferences from Ancient DNA of Pigs and Cattle. Scientific Reports, 7(1).

Meiri, Meirav, Huchon, D., Bar-Oz, G., Boaretto, E., Horwitz, L. K., Maeir, A. M., Sapir-Hen, L., Larson, G., Weiner, S., & Finkelstein, I. (2013). Ancient DNA and population turnover in southern levantine pigs--signature of the sea peoples migration? Scientific Reports, 3, 3035.

Muñiz, A. M., Pecharroman, M. A. C., Carrasquilla, F. H., & von Lettow-Vorbeck, C. L. (1995). Of mice and sparrows: Commensal faunas from the Iberian iron age in the duero valley (central spain). International Journal of Osteoarchaeology, 5(2), 127–138.

Nieto-Espinet, A., Valenzuela-Lamas, S., Bosch, D., & Gardeisen, A. (2020). Livestock production, politics and trade: A glimpse from Iron Age and Roman Languedoc. Journal of Archaeological Science: Reports, 30, 102077.

Nitschke, J. L., Martin, S. R., & Shalev, Y. (2011). Between the Carmel and the Sea – Tel Dor: The Late Periods. Near Eastern Archaeology, 74(3), 132–154.

Ottoni, C., Flink, L. G., Evin, A., Geörg, C., De Cupere, B., Van Neer, W., Bartosiewicz, L., Linderholm, A., Barnett, R., Peters, J., Decorte, R., Waelkens, M., Vanderheyden, N., Ricaut, F.-X., Cakirlar, C., Cevik, O., Hoelzel, A. R., Mashkour, M., Karimlu, A. F. M., Larson, G. (2013). Pig domestication and human-mediated dispersal in western Eurasia revealed through ancient DNA and geometric morphometrics. Molecular Biology and Evolution, 30(4), 824–832.

Panitz-Cohen, N., Mullins, R. A., & Bonfil, R. (2013). Northern Exposure: Launching Excavations at Tell Abil el-Qameh (Abel Beth Maacah). Strata: Bulletin of the Anglo-Israel Archaeological Society, 31, 27–42.

Panitz-Cohen, N., & Yahalom-Mack, N. (2019). The Wise Woman of Abel Beth Maacah. Biblical Archaeology Review, 45(4), 26–33.

Phelps, W., Lolos, Y., & Vichos, Y. (Eds.). (1999). The Point Iria Wreck: Interconnections in the Mediterranean, ca. 1200 BC. Helenic Institute of Maritime Archaeology.

Pöllath, N., Schafberg, R., & Peters, J. (2019). Astragalar morphology: Approaching the cultural trajectories of wild and domestic sheep applying Geometric Morphometrics. Journal of Archaeological Science: Reports, 23, 810–821.

Price, M. D., Perry-Gal, L., & Reshef, H. (2023). The Southern Levantine pig from domestication to Romanization: A biometrical approach. Journal of Archaeological Science, 157, 105828.

Pulak, C. (1998). The Uluburun shipwreck: an overview. International Journal of Nautical Archaeology, 27(3), 188–224.

Raban, A. (1981). Some Archaeological Evidence for Ancient Maritime Activities At Dor. Sefunim, 15–26.

Raban-Gerstel, N., Zohar, I., Bar-Oz, G., Sharon, I., & Gilboa, A. (2008). Early Iron Age Dor: A Faunal Perspective. Bulletin of the American Schools of Oriental Research, 349, 25–59.

Raptou, E. (2008). Kouklia-Lingrin tou Digeni. Report of the Department of Antiquities, Cyprus (RDAC), 2007, 72–73.

Raptou, E. (2010). Kouklia-Lingrin tou Digeni. Report of the Department of Antiquities, Cyprus (RDAC), 2008, 86–87.

Raveh, K., & Kingsley, S. A. (1991). The Status of Dor in Late Antiquity: A Maritime Perspective. The Biblical Archaeologist, 54(4), 198–207.

Robin, O., & Clavel, B. (2018). The diversity evolution of sheep morphology in French zooarchaeological remains from the 9th to the 19th century: Analysis of pastoral strategy. Journal of Archaeological Science, 99, 55–65.

Rohlf, F. J. (2015). The tps series of software. Hystrix, 26(1), 9–12.

Rohlf, F. J. (2017). tpsUTIL, tps file utility program, version 1.74. Stony Brook, NY: Department of Ecology and Evolution, State University of New York at Stony Brook.

Sapir-Hen, L., Bar-Oz, G., Sharon, I., Gilboa, A., & Dayan, T. (2014). Food, Economy, and Culture at Tel Dor, Israel: A Diachronic Study of Faunal Remains from 15 Centuries of Occupation. Bulletin of the American Schools of Oriental Research, 371, 83–101.

Schlager, S. (2017). Morpho and Rvcg -- Shape Analysis in R. In G. Zheng, S. Li, & G. Szekely (Eds.), Statistical Shape and Deformation Analysis (pp. 217–256). xAcademic Press.

Slim, F. G., & Çakirlar, C. (2023). Pigs and polities in Iron Age and Roman Anatolia: An interregional zooarchaeological analysis. Quaternary International: The Journal of the International Union for Quaternary Research, 662-663, 47–62.

Stern, E., Lewinson-Gilboa, A., & Aviram, J. (Eds.). (1993a). The New Encyclopedia of Archaeological excavations in the Holy Land: Volume 2. Jerusalem: The Israel Exploration Society & Carta. 10.5860/choice.31-1294

Stern, E., Lewinson-Gilboa, A., & Aviram, J. (Eds.). (1993b). The New Encyclopedia of Archaeological Excavations in the Holy Land: Volume 4. Jerusalem: The Israel Exploration Society & Carta.

Susnow, M., Marom, N., & Shatil, A. (2021). Contextualizing an Iron Age IIA Hoard of Astragali from Tel Abel Beth Maacah, Israel. Journal of Mediterranean Archaeology, 34(1), 58–83.

Todd, N. B. (1977). Cats and Commerce. Scientific American, 237(5), 100–107.

Trentacoste, A., Nieto-Espinet, A., & Valenzuela-Lamas, S. (2018). Pre-Roman improvements to agricultural production: Evidence from livestock husbandry in late prehistoric Italy. PloS One, 13(12), e0208109.

Valenzuela-Lamas, S., & Albarella, U. (2017). Animal Husbandry across the Western Roman Empire: Changes and Continuities. European Journal of Archaeology, 20(3), 402–415.

Valenzuela-Lamas, S., Orengo, H. A., Bosch, D., Pellegrini, M., Halstead, P., Nieto-Espinet, A., Trentacoste, A., Jiménez-Manchón, S., López-Reyes, D., & Jornet-Niella, R. (2018). Shipping amphorae and shipping sheep? Livestock mobility in the north-east Iberian peninsula during the Iron Age based on strontium isotopic analyses of sheep and goat tooth enamel. PloS One, 13(10), e0205283.

Vigne, J.-D., Zazzo, A., Cucchi, T., Carrère, I., Briois, F., & Guilaine, J. (2014). The transportation of mammals to Cyprus sheds light on early voyaging and boats in the Mediterranean Sea. Eurasian Prehistory, 10(1-2), 157–176

Vuillien, M. (2020). Systèmes d’élevage et pastoralisme en Provence et dans les Alpes méridionales durant la Protohistoire: Nouvelles perspectives en archéozoologie [Doctoral dissertation]. Université Côte d’Azur.

Wachsmann, S., & Raveh, K. (1984). A concise nautical history of Dor/Tantura. International Journal of Nautical Archaeology, 13(3), 223–241. 10.1111/j.1095-9270.1984.tb01194.x

Wickham, H., Averick, M., Bryan, J., Chang, W., McGowan, L. D., François, R., Grolemund, G., Hayes, A., Henry, L., Hester, J., Kuhn, M., Pedersen, T. L., Miller, E., Bache, S. M., Müller, K., Ooms, J., Robinson, D., Seidel, D. P., Spinu, V., … Yutani, H. (2019). Welcome to the tidyverse. In Journal of Open Source Software, 4(43).

Xiao, N. (2018). ggsci: Scientific Journal and Sci-Fi Themed Color Palettes for “ggplot2.” https://CRAN.R-project.org/package=ggsci

Yahalom-Mack, N., Panitz-Cohen, N., & Mullins, R. (2018). From a Fortified Canaanite City-State to “a City and a Mother” in Israel: Five Seasons of Excavation at Tel Abel Beth Maacah. Near Eastern Archaeology, 81(2), 145–156.

Yasur-Landau, A., Shalev, E. A., Zajac, P. R., & Gambash, G. (2018). Rethinking the Anchorages and Harbours of the Southern Levant 2000 BC - 600 AD. In C. von Carnap-Bornheim, F. Daim, P. Ettel, & U. Warnke (Eds.), Harbours as objects of interdisciplinary research – Archaeology + History + Geosciences (pp. 73–90). Mainz: Verlag des Römisch-Germanischen Zentralmuseums.

Yezerinac, S. M., Lougheed, S. C., & Handford, P. (1992). Measurement Error and Morphometric Studies: Statistical Power and Observer Experience. Systematic Biology, 41(4), 471–482.

Zeder, M. A., & Lapham, H. A. (2010). Assessing the reliability of criteria used to identify postcranial bones in sheep, Ovis, and goats, Capra. Journal of Archaeological Science, 37(11), 2887–2905.

Zelditch, M. L., Swiderski, D. L., & Sheets, H. D. (2014). Geometric Morphometrics for Biologists: A Primer (2nd ed.). Academic Press.

